# Sickle red blood cell derived extracellular vesicles activate endothelial cells and enhance sickle red cell adhesion mediated by von Willebrand factor

**DOI:** 10.1101/2022.05.25.492883

**Authors:** Ran An, Yuncheng Man, Kevin Cheng, Tianyi Zhang, Chunsheng Chen, Erdem Kucukal, William J. Wulftange, Utku Goreke, Allison Bode, Lalitha V. Nayak, Gregory M. Vercellotti, John D. Belcher, Jane A. Little, Umut A. Gurkan

## Abstract

Endothelial activation and sickle red blood cell (RBC) adhesion are central to the pathogenesis of sickle cell disease (SCD). Quantitatively, RBC-derived extracellular vesicles, REVs, are more abundant from SS RBCs compared with healthy RBCs (AA RBCs). Sickle RBC-derived REVs (SS REVs) are known to promote endothelial cell (EC) activation through cell signaling and transcriptional regulation at longer terms. However, the SS REV-mediated short term non transcriptional response of EC is unclear. Here, we examined the impact of SS REVs on acute microvascular EC activation and RBC adhesion at 2 hours. Compared with AA REVs, SS REVs promoted human pulmonary microvascular endothelial cells (HPMEC) activation indicated by increased von Willebrand Factor (vWF) expression. Under microfluidic conditions, we found abnormal SS RBC adhesion to HPMECs exposed to SS REVs. This enhanced SS RBC adhesion was reduced by vWF cleaving protease ADAMTS13 to a level similar to HPMECs treated with AA REVs. Consistent with these observations, studies in SS mice with implanted dorsal skin-fold chambers found hemin-induced stasis was inhibited by ADAMTS13. The adhesion induced by SS REVs was variable, and was higher with SS RBCs from patients with increased markers of hemolysis (LDH and reticulocyte count) or a concomitant clinical diagnosis of deep vein thrombosis. Our results emphasize the critical contribution made by REVs to the pathophysiology of SCD by triggering acute microvascular EC activation and abnormal RBC adhesion. These findings may help to better understand acute pathophysiological mechanism of SCD and thereby the development of new treatment strategies using vWF as a potential target.

## 1. INTRODUCTION

Sickle cell disease (SCD) is a genetically inherited blood disorder, in which a point mutation in the beta globin chain gene replaces A with T at codon 6. This causes a switch from a hydrophilic glutamic acid to a hydrophobic valine, producing abnormal sickle hemoglobin (HbS). HbS polymerizes into long and stiff intracellular structures under dehydration and deoxygenation, leading to sickle-shaped, excessively stiff, and adhesive red blood cells (RBCs). Sickle RBCs (SS RBCs) are prone to extra- and intravascular hemolysis [1]. During hemolysis, SS RBCs release hemoglobin (Hb), heme, and extracellular vesicles (EVs) [2–6], which cumulatively promote a pro-inflammatory milieu and consequently intermittent vasoocclusion of the microvasculature in SCD [2, 7].

EVs are membrane-bound sub-micron particles (0.1-1.0 μm) generated from cells under normal and activated conditions [8]. EVs contain various cargos including proteins, lipids, and nucleic acids, which may reflect the state of activation of the cells from which they originate [9] and can serve as vehicles for cellular communication [8]. The level of total circulating EVs (including EVs from platelets, leukocytes, endothelial cells, and RBCs) have been reported to be higher in the bloodstream of subjects with SCD at steady state than in healthy controls and increased during SCD vaso-occlusive crises [10, 11]. RBC-derived EVs (REVs) made up the most prevalent subtype [12], and correlated with hemolysis, oxygen saturation, pulmonary systolic pressure, and mortality [13]. REVs in SCD (SS REVs) may be produced through accelerated aging of RBCs during oxygenation/deoxygenation cycles [16], under the influence of stress factors [4, 14–16], or during hemolysis in capillary beds. SS REVs are reported to have increased phosphatidylserine (PS) expression on their surface [17], as well as to contain heme, Hb, oxidized Hb, ferryl Hb, and microRNAs [3, 18, 19]. SS REVs are known to have a range of pathophysiologic impacts on blood-vascular interactions including nitric oxide scavenging [20], immune modulation [21], coagulation [22], decrease endothelial monolayer integrity [23–25], and adhesion of blood cells to the endothelium [5, 23]. Garnier et al. showed SS REV-treated ECs upregulated mRNA and protein expression of intercellular adhesion molecule-1 (ICAM-1) at longer terms (4 or 6 hours), which led to significantly enhanced neutrophil recruitment to human endothelium in SCD [5]. Gemel and Lapping-Carr et al. reported that REVs in plasma EVs isolated from subjects with SCD during acute chest syndrome significantly decreased levels of endothelial VE-cadherin protein and disrupted junctions at 48 hours [24, 25]. Camus et al. demonstrated that in transgenic SAD mice, SS REVs are capable of transferring heme to endothelial cells (ECs), triggering microvascular-occlusions, likely through toll-like receptor-4 (TLR-4) dependent pathways [3]. Importantly, much of the cell-free heme in plasma is bound to REVs [3] therefore endothelial activation *via* cell-free heme is likely mediated by the transfer of heme through REVs [3]. However, most of the existing work indicate SS REVs promote endothelial activation at longer terms (>4 hours) through transcriptional responses, while the acute (≤2 hours) impact of REVs on endothelial activation has not been fully elucidated.

In this study, we utilized our previously developed endothelium-on-a-chip platform [26] to analyze RBC adhesion to REV-activated human pulmonary microvascular endothelial cells (HPMECs) under physiologic flow conditions. SS REVs were generated by increasing the intracellular calcium concentration in SS RBCs. Following short-term (2 hours) REV activation of microvascular ECs, which mimics the proinflammatory phenotype of microvascular endothelium during acute hemolysis in SCD, we analyzed RBC adhesion to the HPMECs using clinical blood samples from subjects with homogeneous SCD followed in our clinic. Here, we report SS RBC adhesion on REV-activated HPMECs and its clinical associations in SCD. The aim of the present study is to demonstrate the acute impact of REVs generated from SS RBCs on microvascular EC phenotype, focusing on abnormal RBC adhesion.

## 2. MATERIALS AND METHODS

### 2.1. Blood sample collection

All experiments were performed in accordance with approved study protocol by the Institutional Review Board (IRB) committee (IRB 05-14-07C). Blood samples from de-identified adult subjects with HbSS or healthy donors (HbAA) were obtained at University Hospitals Cleveland Medical Center (UHCMC) in Cleveland, Ohio. Hb composition of SCD samples was identified by high-performance liquid chromatography (HPLC) at the Core Laboratory of UHCMC. Clinical variables of the study subjects with SCD were obtained from the medical record.

### 2.2. Microfluidic channel endothelialization

Microfluidic channels were fabricated as previously described [34–38]. Briefly, the microchannels were fabricated by assembling three layers, including a glass slide as bottom layer, double-sided adhesive (DSA) film as middle layer, and polymethyl methacrylate (PMMA) as top layer. Microchannel geometry was determined by the middle DSA layer and channel inlet and outlet was determined on the top PMMA layer (**Fig. 1**). The microchannels were incubated with 0.2 mg/mL fibronectin for 1 hour at 37°C. HPMECs were seeded into the microchannels and cultured with 5% CO_2_ at 37°C under 100 μL/min flow for 48-72 hours until a confluent monolayer over the microchannel bottom surface was formed (**Fig. 1**).

**Figure 1.**
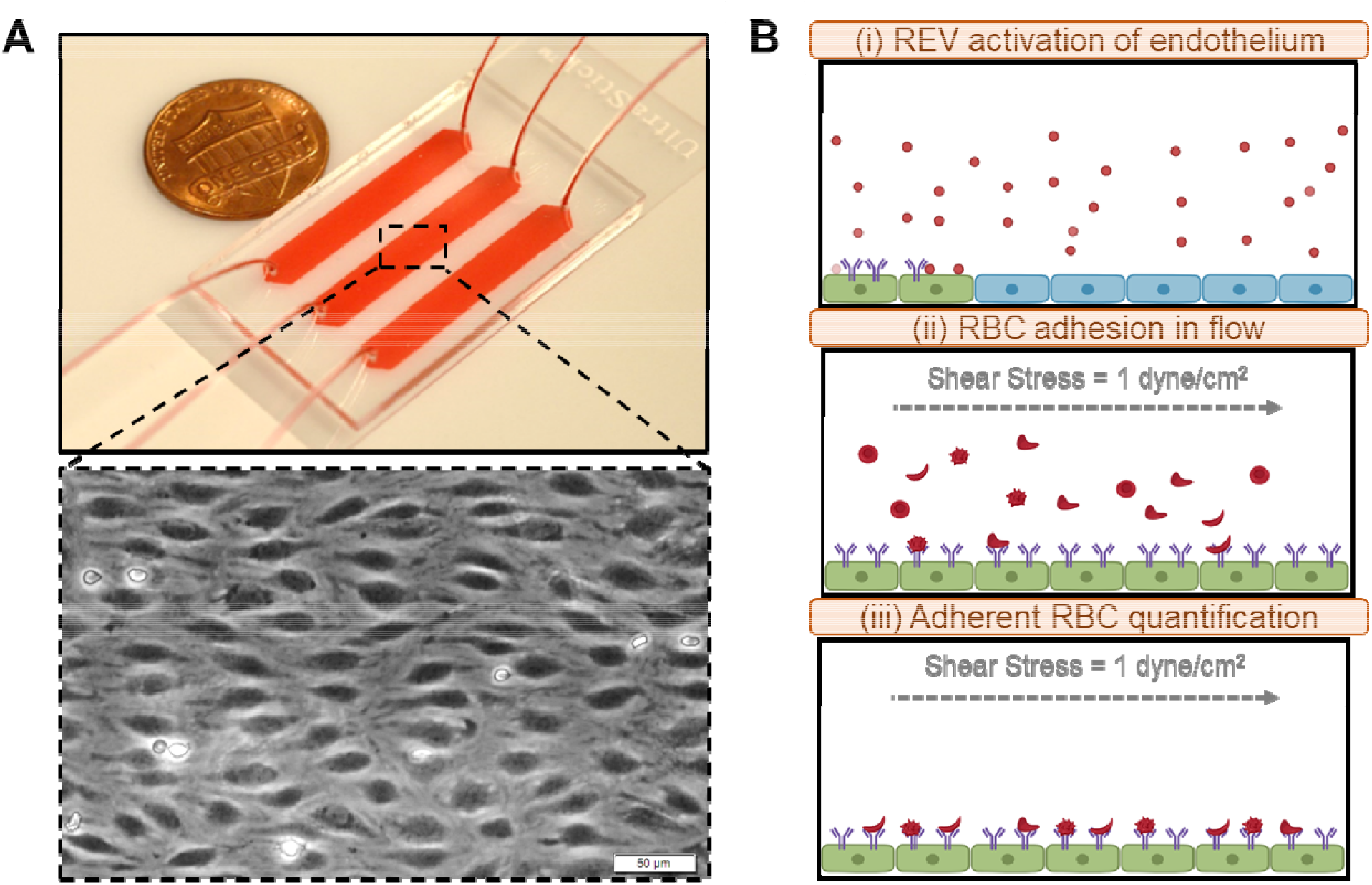
The endothelium-on-a-chip system for probing RBC adhesion to REV activated endothelial cells: **(A)** Overview of the endothelium-on-a-chip system with a clinical SCD sample flown over HPMECs. (B) Overview of the microfluidic assay test procedure: (i) In vitro generated REVs are firstly perfused into endothelialized microchannels; (ii) Upon treatment, 15 μL re-suspended RBC solutions from blood samples in individual patients were perfused into the microchannels at physiologically relevant shear stress of 1 dyne/cm2, for a; and (iii) non-adherent RBCs were washed off by basal medium and the adherent RBCs were quantified.

### 2.3. Red blood cell derived extracellular vesicle (REV) generation and characterization

Pooled blood samples from 10 subjects with HbSS and 5 healthy donors were used to derive HbSS REV and HbAA REV, respectively. RBCs were isolated from either individual or pooled blood samples by centrifuging for 5 minutes at 500×g at room temperature. Plasma, buffy coat, and the near-plasma portion of the RBC layer were carefully removed. The isolated RBCs were washed twice with PBS, and were resuspended in Hank’s balanced salt solution modified with calcium and magnesium (Hank’s buffer) at 20% hematocrit. The RBC suspensions were then stimulated with 2 μM calcium ionophore (A23187) for 1 hour at 37°C. To isolate generated REVs, cell suspensions were subject to differential centrifugation, initially at 1,500 ×g for 15 min followed by the second-step centrifugation at 3000×g for 15 min. Finally, the supernatants were centrifuged at 25,000×g for 2 hours at 4°C to harvest REV pellets. REVs were resuspended in either cell culture media (for endothelial cell activation) or PBS (for REV characterization), split into 200 μL aliquots and stored in −80°C before using. The size distribution and concentration of the generated REVs were measured using ZetaView^®^ particle tracking analyzer (Particle-Metrix, Germany).

### 2.4. Endothelial cell activation

For all endothelial activation experiments without hemopexin, stock REVs were thawed at 4 °C and brought to 37 °C in the cell culture incubator. To test the impact of hemopexin on REV-mediated HPMEC activation, REVs were incubated with 2 μM human hemopexin (Sigma-Aldrich) for 1 hour at room temperature. For all experiments, cultured HPMECs (LONZA) were washed with fresh culture medium and were then incubated with HbSS REVs or HbAA REVs for 2 hours at 37°C. The activated HPMECs were then washed with fresh culture medium and used for adhesion experiments. In control experiments, SS REVs and AA REVs were reconstituted to the same concentration.

### 2.5. Fluorescent labeling of von Willebrand factor (vWF) on endothelial cells

Following REV activation, HPMECs were rinsed with fresh culture medium and fixed with 4% paraformaldehyde (PFA) for 15 minutes at room temperature. Fixed HPMECs were rinsed with PBS twice and blocked with 2% BSA for 1-hour at room temperature. After washing with PBS, HPMECs were incubated with sheep polyclonal anti-human vWF antibody (Abcam) conjugated with fluorescein isothiocyanate (FITC, 1:100 v/v dilution) for 1 hour at room temperature in dark. Fluorescent images were then acquired at multiple locations throughout the microchannel of REV activated HPMECs at 10X.

### 2.6. Adhesion Experiments

For all RBC adhesion experiments without VWF cleaving protease ADAMTS13, RBCs were isolated from individual blood samples by centrifugation for 5 minutes at 500×g at room temperature. The isolated RBCs were washed twice with PBS and were re-suspended in fresh basal medium supplemented with 10 mM HEPES at 20% hematocrit. To test the impact of ADAMTS13 on RBC adhesion, freshly isolated RBCs in basal medium supplemented with 10 mM HEPES at 20% hematocrit were incubated for 30 minutes under 37 °C with gentle shaking with 5 μg/mL ADAMTS13 (R&D Systems) [27, 28].

For all RBC adhesion experiment, a total sample volume of 15 μL was perfused into the microchannel at the shear rate of 1 dyne/cm^2^, corresponding to a typical value observed in human post-capillary venules. After blood perfusion, the microchannels were rinsed with fresh basal medium supplemented with 10 mM HEPES to remove non-adherent RBCs. Visual confirmation of adhesion results confirmed negligible contamination by platelets or white blood cells.

### 2.7. Mice

All animal experiments were approved by the University of Minnesota’s Institutional Animal Care and Use Committee. In this study, we utilized male and female HbSS-Townes on a 129/B6 mixed genetic background. The SS mice were created by knocking in human α and ^A^γβ^S^ globins into the sites where murine α-globin and β-globins were knocked out [29]. SS mice have severe anemia and an SS RBC halflife of 2.5 days [29]. All animals were monitored daily including weekends and holidays for health problems, food and water levels and cage conditions. Littermates were randomly assigned to different treatment groups. All animals were included in each endpoint analysis and there were no unexpected adverse events that required modification of the protocol. Mice were aged 11-14 weeks.

### 2.8. Statistical Methods

Mean ± standard deviation (SD) of the mean were reported for acquired data in this study. Minitab 18 Software (Minitab Inc., State College, PA) was used to perform all statistical analyses. Data normality was initially analyzed. For comparison, normally distributed data were analyzed by parametric one-way ANOVA, and non-normally distributed data were analyzed by non-parametric Mann-Whitney U test. Statistical significance was set at 95% confidence level for all tests (P < 0.05).

## 3. RESULTS

### 3.1. Characterization of red blood cell derived extracellular vesicles (REVs)

We characterized and compared calcium ionophore-derived AA REVs, generated from the pooled blood samples from 5 healthy donors, and calcium ionophore-derived SS REVs, generated from the pooled blood samples from 5 people with SCD (**Fig. 2**). The nanoparticle tracking analysis (NTA) results demonstrated that derived AA and SS REVs had similar diameters (10 to 700 nm with >95% of them between 80-400 nm, **Fig. 2A**, mean size ± SD: AA REV = 205.4 ± 223.7 nm vs. SS REV = 181.7 ± 200.2 nm). However, per RBC volume, SS RBCs generated significantly more REVs than did AA RBCs (**Fig. 2B**, mean concentration ± SD: SS REV = 3.17 × 10^7^ ± 3.91 × 10^6^/mL vs. AA REV = 6.35 × 10^6^ ± 5.68 × 10^5^/mL, p = 0.011, Mann-Whitney).

**Figure 2.**
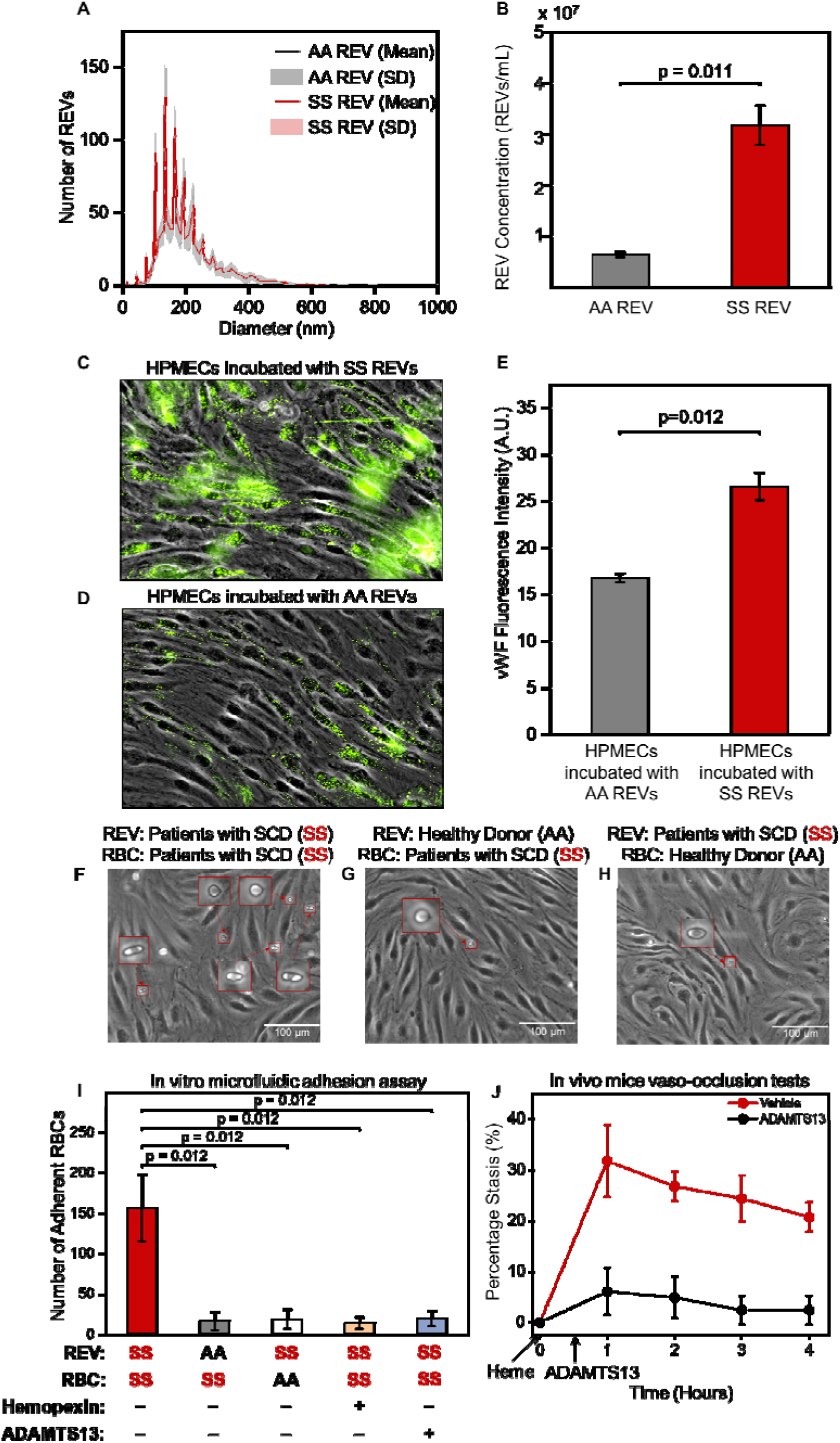
SS RBCs generate higher levels of REVs which activate HPMECs, increase vWF expression and mediates enhanced adhesion of SS RBCs, which can be reduced by hemopexin and ADAMTS13. **(A)** Size distribution of AA REVs and SS REVs generated from RBCs in pooled-samples from healthy donors or pooled-samples from patients with SCD. Black and red lines indicate the mean number of AA and SS REVs from 5 individual tests and gray and light-red shaded area indicate the standard deviation. AA REV and SS REV share similar size distribution (Mean size ± SD: AA REV = 205.4 ± 223.7 nm vs. SS REV = 181.7 ± 200.2 nm). **(B)** Concentration of AA REV and SS REV generated in vitro per microliter of purified RBCs in samples from either healthy donors (gray) and patients with SCD (red). SS RBCs generated statistically significantly higher quantity of REVs than AA RBCs (Mean concentration ± SD: SS REV = 3.17 × 10^7^ ± 3.91 × 10^6^/mL vs. AA REV = 6.35 × 10^6^ ± 5.68 × 10^5^/mL, p = 0.011, Mann-Whitney). **(C, D)**: HPMECs were treated with SS REVs (**C**) and AA REVs (**D**) for 2 hours in 37 °C and incubated with fluorescently labeled antibodies against vWF following a fixing step with 4% PFA. The SS REV treated HPMECs demonstrated significantly increased vWF expression comparing to HPMECs incubated with AA REVs **(E,** n=5, p=0.012, Mann–Whitney U test). **(F)** SS RBC adhesion on HPMECs activated with SS REVs. **(G)** SS RBCs adhesion on HPMECs treated with AA REVs. **(H)** AA RBCs adhesion on HPMECs treated with SS REVs. Insets are closer view of RBCs adhered to HPMECs. **(I)** Interaction between SS RBC and SS REV activated HPMECs is significantly stronger than the other four test groups, and this interaction is reduced by hemopexin and ADAMTS13 (mean ± SD: SS REV-SS RBC = 157 ± 42, AA REV-SS RBC = 16 ± 12, AA REV-SS RBC = 19 ± 12, SS REV-SS RBC with hemopexin = 14 ± 8, SS REV-SS RBC with ADAMTS13 = 19 ± 10, p < 0.05 for all groups, n = 5 in each group, Mann–Whitney U test). **(J)** Heme-induced vaso-occlusion (measured in percentage of stasis) in vivo in SS Towns mice is inhibited by ADAMTS13 (mean ± SD for 1, 2, 3, and 4 hours: Vehicle: 31.8% ± 7.1%, 26.8% ± 2.9%, 24.5% ± 4.5%, 20.8% ± 2.9% vs. ADAMTS13: 6.1% ± 4.7%, 4.9% ± 4.1%, 2.4% ± 2.8%, 2.4% ± 2.8%, p < 0.05 for all time points, n = 4 for each group at each time point)

### 3.2. SS REVs increase endothelial von Willebrand factor (vWF) expression

We incubated HPMECs with derived SS or AA REVs for 2 hours. vWF expression was significantly increased on HPMECs incubated with derived SS REVs (**Fig. 2C, D**) compared to vWF expression on HPMECs incubated with derived AA REVs (**Fig. 2E**, p = 0.012, Mann-Whitney).

### 3.3. HbSS RBC adhere specifically to HPMECs incubated with HbSS REV

We first verified the in vitro approach by examining the interactions between SS RBC, AA RBC, and HPMECs treated with SS REVs and AA REVs. HPMEC laminated microchannels were treated with derived SS REV (**Fig. 2F and 2H**) or derived AA REV (**Fig. 2G**). SS RBC suspensions were then perfused into the first 2 channels (**Fig 2F & G**), and AA RBC suspension was perfused into the third (**Fig 2H,** per **Section 2.5)**. No visible injury or morphological changes were observed on HPMECs treated with SS REV or AA REV. More adherent SS RBCs were observed on HPMECs activated with SS REVs than with AA REVs (**Fig. 2I**, 157 ± 42 vs. 19 ± 12, p < 0.05) and AA RBC on HPMECs treated with SS REVs (**Fig. 2I**, 16 ± 12, p < 0.05).

### 3.4. REV-mediated RBC adhesion to HPMECs are attenuated by hemopexin

To demonstrate that the observed RBC adhesion was induced by REV-mediated HPMEC activation, we investigated the impact of the heme-binding protein hemopexin on REV-mediated RBC adhesion. Pre-incubating REVs with hemopexin reduced the HPMEC activation indicated by reduced number of adherent SS RBCs (**Fig. 2I,** 157 ± 42 vs. 14 ± 8, p < 0.05). These results indicate that REV-mediated HPMEC activation was dependent on heme content, because hemopexin dramatically inhibited HPMEC activation, indicated by reduced number of RBC adhesion.

### 3.5. REV-mediated RBC adhesion to HPMECs are attenuated by ADAMTS13

To demonstrate that the observed RBC adhesion were mediated by vWF on REV-activated HPMECs, we investigated the impact of the vWF-specific protease ADAMTS13 on REV-mediated RBC adhesion. The addition of ADAMTS13 significantly reduced the adhesion of SS RBCs to SS REV-activated HPMECs (**Fig. 2I,** 157 ± 42 vs. 19 ± 10, p < 0.05). These results indicate that RBC adhesion was dependent on HPMEC vWF, because ADAMTS13 dramatically inhibited RBC adhesion, reaching a level that was close to SS RBC on HPMECs treated with AA REVs.

### 3.6. Heme-induced vaso-occlusion in SS Towns mice is inhibited by ADAMTS13

Since ADAMTS13 reduced RBC-endothelial cell adhesion and RBC adhesion to endothelium is believed to be involved in vaso-occlusion in SCD, we used Townes SS mice and a dorsal skin-fold chamber model to determine if ADAMTS13 would reduce microvascular stasis in response to hemin. Townes SS mice were surgically implanted with a dorsal skin-fold chamber. After chamber implantation, 20-21 subcutaneous flowing venules were selected at baseline using intravital microscopy. After selection and mapping of flowing venules, SS mice were infused via the tail vein with hemin (3.2 μmol/kg) at time zero to induce vaso-occlusion [3]. Twenty minutes later, mice were infused with ADAMTS13 (1 mg/kg) or sterile saline (10 ml/kg). The same venules selected at baseline were re-examined for stasis (no flow) at 1, 2, 3, and 4 hours after hemin infusion. Vaso-occlusion (stasis) was significantly inhibited at 1, 2, 3, and 4 hours post-hemin in SS mice that received ADAMTS13 compared to mice that received saline (**Fig. 2J**). These results demonstrate that the ADAMTS13 protease can disrupt vaso-occlusion induced by heme, which is the main component in REVs.

### 3.7. Clinical implications of REV-mediated RBC adhesion to HPMECs

To determine the association between clinical phenotypes and adhesion profiles, we examined a hemolytic subpopulation (Group 1, N = 6) with distinctly lower lactate dehydrogenase (LDH) levels and absolute reticulocyte counts (ARCs) compared with the 2^nd^ subpopulation (Group 2, N = 9, **Fig. 3A**) [42]. Subjects in Group 1 had significantly lower LDH levels and ARCs (N = 8, LDH mean ± standard deviation: Group 1: 186 ± 99 IU vs. Group 2: 305 ± 55 IU, p < 0.034, Mann-Whitney; ARCs mean ± standard deviation: Group 1: 162 ± 74 vs. Group 2: 401 ± 122 × 10^10^, p < 0.001, One-way ANOVA). We then compared the RBC adhesion profile between Group 1 subjects and Group 2 subjects, and found significantly lower RBC adhesion to HbS REV activated HPMECs in Group 1 subjects than in Group 2 subjects (**Fig. 3B**, 117 ± 72 vs. 267± 125, p = 0.016, Mann-Whitney). Subjects in Group 1 also had lower WBCs than did subjects in group 2 (**Fig. 3C**, 8.5 ± 1.2 × 10^9^ vs. 11.4 ± 3.6 × 10^9^ / L, p = 0.034, Mann-Whitney). Group 1 also had a suggestion of having lower ferritin values (**Fig. 3D**, 876 ± 1527 ug/L vs. 2841 ± 2782 μg/L, p = 0.057, Mann-Whitney).

**Figure 3.**
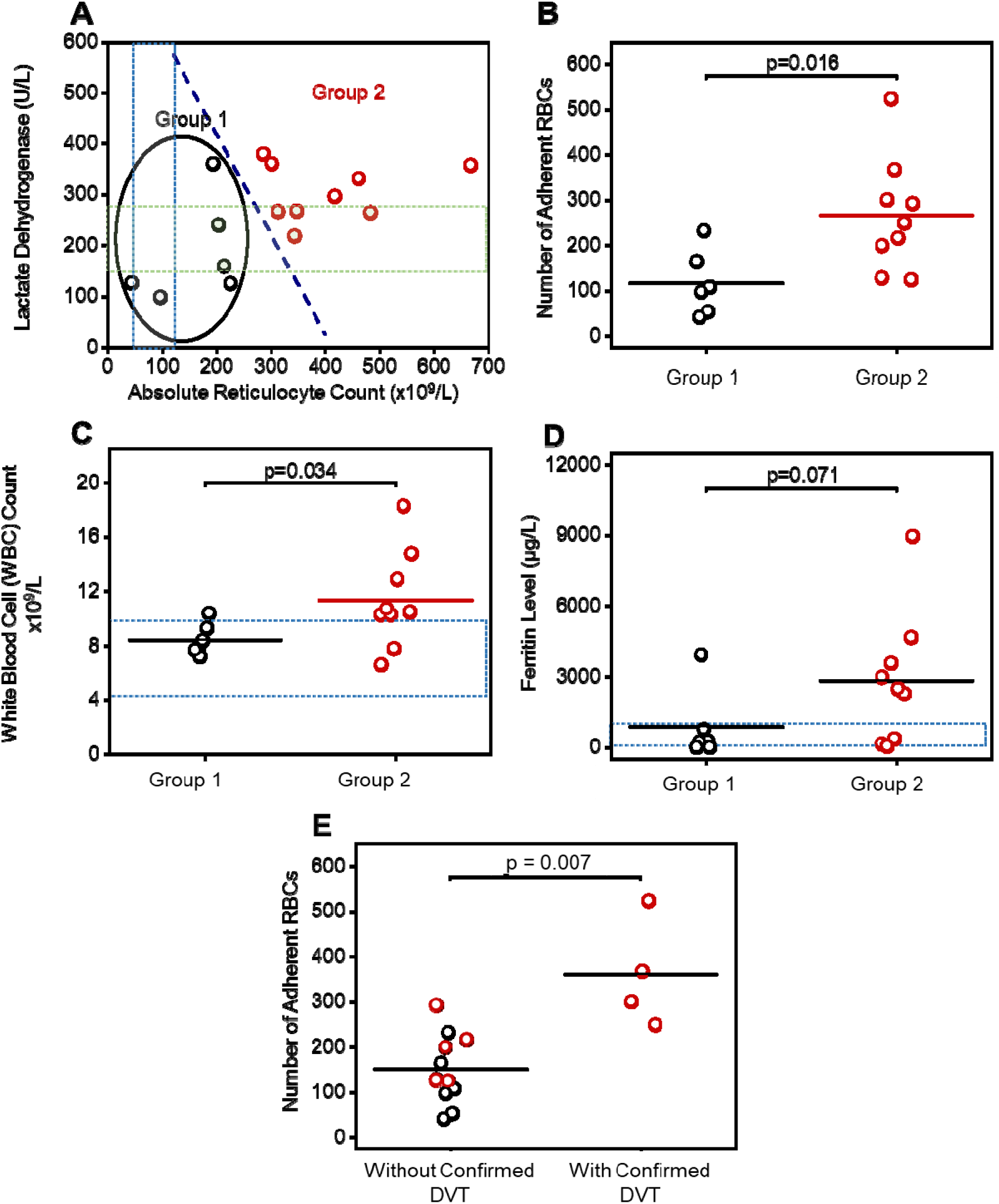
RBC adhesion to REV-activated HPMECs correlates with subject clinical phenotype including hemolytic and inflammatory biomarkers. **(A)**: A subpopulation (Group 1, N = 6) with distinct hemolysis markers of lactate dehydrogenase (LDH) levels and absolute reticulocyte count (ARCs) comparing to the rest (Group 2, N = 9) via k-means clustering analysis. RBCs from subjects in Group 2, with significantly higher LDH levels and ARCs, have greater adhesion to REV-activated HPMECs compared to the RBCs from subjects in Group 1 (**B**, Mean ± SD: 267 ± 125 vs. 117 ± 72, p = 0.016,One-way ANOVA). The gray and green shaded areas indicate normal ranges for ARC and LDH, respectively. (**C**): Subjects in Group 2 with higher LDH and ARC and enhanced RBC adhesion have significantly higher WBC counts, than subjects in Group 1 (Mean ± SD: 11.4 ± 3.6 vs. 8.5 ± 1.2, p = 0.034, one-tailed Mann-Whitney). Shadowed area: WBC count range from 4.5 to 10 × 10^9^. (**D**): Subjects in Group 2 with higher LDH and ARC and enhanced RBC adhesion have higher ferritin levels, although not statistically significant, than subjects in Group 1 (2841 ± 2804 vs. 876 ± 1527, p = 0.071, one-tailed Mann-Whitney). Shadowed area: normal range of ferritin level (2 to 1000 μg/L) [71, 72]. **(E)** Subjects with confirmed deep vein thrombosis (DVT) had significantly higher RBC adhesion to REV activated HPMECs than the ones without confirmed DVT (Mean ± SD: 361 ± 119 vs. 151 ± 78, p = 0.007, Mann-Whitney). Six out of six subjects in Group 1 patient with lower RBC adhesion, and three out of six subjects in Group 2 did not have DVT. Four out of eight subjects in group 2 with higher RBC adhesion had DVT. DVT status of two out of eight subjects in Group 2 were not available (1 patient diagnosed as ‘unclear’, 1 patient record not accessible).

Finally, regardless of interest, subjects with a documented history of deep vein thromboses (DVT, N = 4) had higher RBC adhesion to REV-activated HPMECs compared to subjects without confirmed DVT (N = 11, **Fig. 3E**, 361 ± 119 vs. 151 ± 78, p = 0.007, Mann-Whitneye). Zero out 6 Group 1 subjects and 4 out 7 Group 2 subjects (with definitive DVT record) were diagnosed with DVT. DVT record of 2 subjects in Group 2 were not available (DVT diagnosis for one subject was not clear and DVT record was not accessible for the other subject). Similarly, Group 2 subjects with definitive DVT record had significantly greater possibility of having DVT than subjects in Group 1 (**Table S1**).

## DISCUSSION

Two dominant pathophysiological events have been characterized with SCD: abnormal RBC adhesion and intravascular hemolysis. Intravascular hemolysis promotes endothelial activation and causes endothelial dysfunction; and SS RBCs are known to have reduced deformability [30, 31] and increased adhesivity to the activated endothelium, leading to vaso-occlusion. These mechanisms likely intersect and contribute to acute painful episodes and chronic organ damage [32].

SCD RBC membrane abnormalities include aberrant timing or abnormal persistence during maturation, and abnormal activation by ‘stress signals’, of surface molecules such as VLA-4, CD36, LW and BCAM/LU [33–38]. Cumulative oxidative damage, resulting in excessive PS externalization on the SCD RBC membrane, causes abnormal adhesion [39, 40]. RBC adhesion to activated endothelium has been associated with elevated endothelium adhesion molecules including P-selectin [41], ICAM-1 [42], Vascular Cell adhesion molecule-1 (VCAM-1) [43], E-selectin [44], and vWF [45], as well as subendothelial proteins such as laminin (LN) under various physiological conditions. These abnormal interactions between SCD RBC and activated endothelium plays a pivotal role in the initiation and propagation of unpredictable VOC episodes, thereby contributing to the pathophysiology of SCD.

Intravascular hemolysis releases free heme, Hb, and REVs. Heme induces endothelial activation through TLR-4, resulting in increased expression of VCAM-1, ICAM-1, E-selectin, P-selectin, IL-1, IL-6, IL-8, and tissue factor via a pathway involving the activation of NF-kB phospho-p65 which are known to mediate RBC adhesion [2, 42, 44]. Importantly, up to one-third of cell-free heme in plasma is sequestered in circulating REVs [46]. Therefore, we set out to reproduce this important physiological mechanism in vitro. SS REVs can transfer heme to ECs in annexin-a5-sensitive fashion and have been reported to trigger transcriptional responses from middle- to longer-time periods (4-48 hours) [5, 24, 25, 46, 47] causing endothelial injury, linking hemolysis to chronic vascular injury in SCD mouse [4, 46, 47]. In addition to these long term transcriptional responses, heme is also known to activate acute stress through sentinel pathway mediated by TLR-4 signaling that leads to production of ROS, triggers WPB degranulation and rapid release of vWF to the cell surface, which eventually causes vaso-occlusion [2, 48]. It has been shown that vWF multimers released from endothelial cells results in a significant increase in adhesion of SS RBCs to endothelium *ex vivo* [49], suggesting an important role in the pathophysiology of sickle cell-induced microvascular occlusion [45, 49]. However, the relationship of this finding and SS REVs has not been well-described.

Leveraging the Endothelium-on-a-chip technology, we treated HPMECs with calcium ionophore-generated REVs in order to mimic the highly hemolytic plasma milieu that is often typical in people with HbSS [7]. We demonstrated enhanced vWF expression on HPMECs treated with SS REVs comparing to those treated with AA REVs within 2 hours. This demonstrates that REVs are capable of inducing acute endothelial response in the short term, in addition to known long-term effects. Further, vWF is known to mediate SS RBC adhesion independent of platelets [45, 50], and plasma levels of vWF may correlate with hemolysis in SCD [51]. We found that SS RBCs demonstrate increased adherence to SS REV-activated HPMECs. These adhesion events were dramatically mitigated by heme-binding protein hemopexin or by vWF-specific protease ADAMTS13, reaching levels that were close to SS RBC on HPMECs treated with AA REVs.

Circulating EVs in SCD are known to have pro-thrombotic effect due to their surface tissue factor and PS expression [12]. Circulating REVs in SCD are particularly known to contain significant amounts of cell-free heme and facilitate heme delivery to ECs [5, 52]. Heme is known to rapidly mobilize Weibel-Palade body (WPB) vWF and P-selectin onto EC surface and cause vaso-occlusion SS mice [2]. Here, we demonstrated the effect of hemopexin in mitigating REV-mediated acute endothelial activation. These results agreed with our previous work regarding the effect of hemopexin in mitigating heme-mediated vaso-occlusion in SS mice [2]. Furthermore, in addition to the in vitro experimental results, we confirmed the effect of ADAMTS13 in mitigating heme-mediated vaso-occlusion measured in percent of stasis in vivo using Townes SS mice. Additionally, Belcher et al. previously demonstrated that polyclonal antibodies to vWF blocked stasis induced by heme in SCD mice with dorsal skin-fold chambers [2]. Together, these results suggest a specific interaction between these cellular elements that is potentially mediated by HPMEC vWF expression induced by the heme content in SS REVs (**Fig. 4**). To analyze the heterogeneity of patient RBC adhesion to REV-activated endothelial cells, we uniformly activated HPMECs with SS REVs generated from pooled blood samples. Therefore, the RBC adhesion quantified in this study only depends on the adhesivity of RBCs in individual patients. Accordingly, we describe 2 groups, one of which is characterized by elevated LDH levels and ARCs, and had significantly greater RBC adhesion as compared to the other group (**Fig. 3A&B**). LDH and ARC are important *in vivo* hemolytic biomarkers which have been linked to disease severity [53, 54]. Therefore, these results suggest that RBCs are more adhesive in subjects with a more severe hemolytic phenotype in SCD. Additionally, we found that subjects with a documented history of confirmed deep vein thromboses (DVT, N = 4) had higher RBC adhesion to REV-activated HPMECs compared to subjects without confirmed DVT. We postulate that this is because the higher adhesivity indicates more severe sickling of RBCs, which has been reported to mediate entrapment of these cells within venous clot [55]. In conclusion, in this study, we have improved our Endothelium-on-a-chip approach by activating the endothelial layer with a biologically complex *in vitro* generated REVs obtained from isolated RBC. We believe that this is physiologically relevant in comparison to a simpler biochemical signal such as TNF-α or heme. Our findings indicate that both pathological RBCs with higher adhesivity, and an activated endothelium are required for the observed abnormal interaction. The SS RBCs demonstrate enhanced adhesion to HPMECs activated with SS REVs in a subject specific fashion. This heterogeneity in adhesion solely reflects heterogeneous RBC adhesion to uniformly activated HPMECs (using pooled derived REV). The RBC adhesion profiles associated with hemolytic and inflammatory biomarkers including LDH, ARC, WBC, and ferritin levels. Future investigations will examine RBC adhesion on endothelial cells that are activated with patient-specific REVs, which may better reflect the overall disease process in an individual patient. Understanding the important collective interplay between RBCs and REV-activated microvascular endothelial cells will better characterize the multicellular adhesion paradigm for acute and chronic vaso-occlusion in SCD and may enable us to develop more effective treatment paradigms.

**Figure 4.**
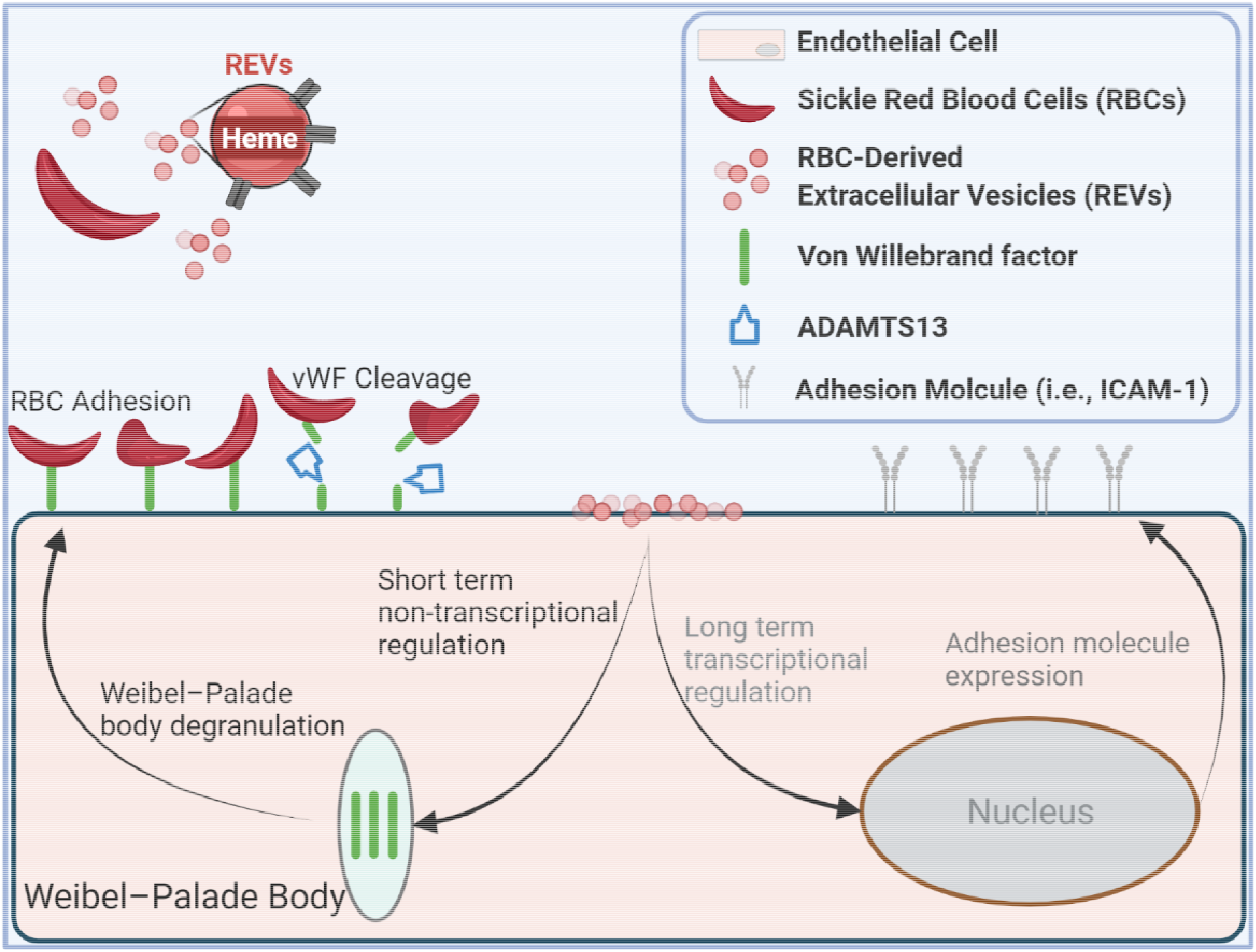
Impact of REV on microvascular endothelial cell response. SS REV are capable of promoting HPMEC VWF expression within 2 hours, likely through the acute stress sentinel pathway of Weibel-Palade body degranulation. SS RBC adhere specifically to SS REV-activated HPMECs, likely mediated by endothelial VWF, and the adhesion is decreased with VWF cleaving protease ADAMTS13. Enhanced SS RBC adhesivity was observed in patients with elevated biomarkers of hemolysis and inflammation and thrombophilia.

## ACKNOWLEDGEMENTS

This work is supported by National Heart, Lung, and Blood Institute (NHLBI) R01HL133574, OT2HL152643, U01HL117659, R01HL114567 (Vercellotti/Belcher), and T32HL134622. This article’s contents are solely the responsibility of the authors and do not necessarily represent the official views of the National Institutes of Health.

## AUTHOR CONTRIBUTIONS

RA and YM contributed equally to this work. RA, YM and UAG conceived the project. RA, YM, KC, CC, and TZ conducted the experiments. RA, YM, TZ, KC, EK, WJW, GMV, JDB, and UG analyzed the data. RA, YM, LVN, JAL, GMV, JDB, and UAG discussed and interpreted the data. RA and YM prepared the figures and table, and wrote the manuscript. RA, YM, LVN, JAL, GMV, JDB, and UAG reviewed and edited the manuscripts. AB collected patient clinical information from the medical record and blood sample collection.

---

**Supplemental Table 1.**
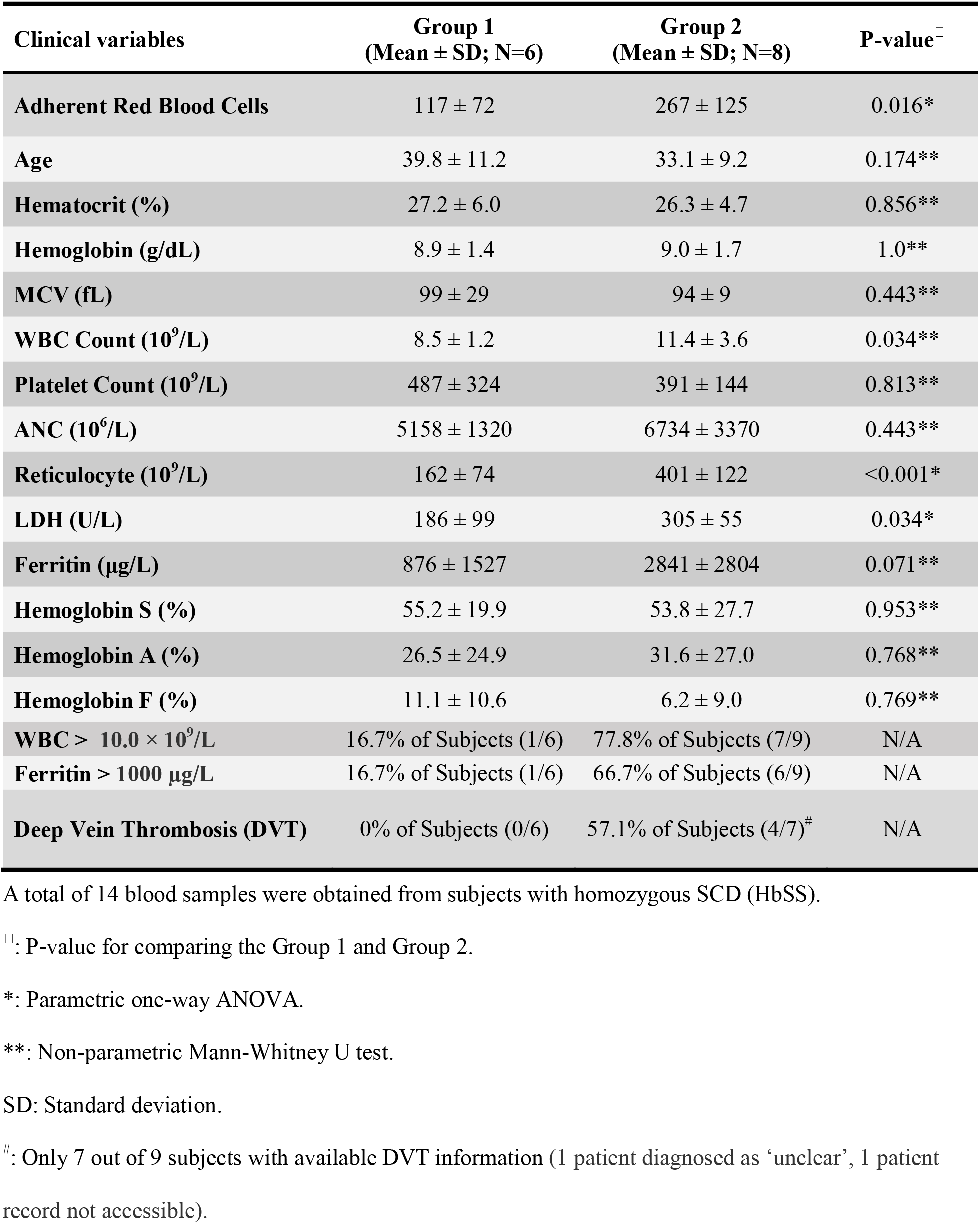
Clinical variables of the study population.

